# Gut Reaction: The Impact of a Film on Public Understanding of Gastrointestinal Conditions

**DOI:** 10.1101/2021.09.05.459031

**Authors:** Diogo Gomes, Alina Loth, James R.F. Hockley, Ewan St. John Smith

**Affiliations:** Department of Pharmacology, University of Cambridge, Tennis Court Road, Cambridge, CB2 1PD

## Abstract

Chronic gastrointestinal (GI) tract conditions, such as inflammatory bowel disease (IBD) and irritable bowel syndrome (IBS) are common conditions associated with disordered bowel movements and significant pain. However, discussion of bowel habits is often regarded as taboo and public understanding of what exactly IBD, IBS and related conditions are, and how they impact the lives of those individuals with such conditions is poorly understood. To provide a platform for enhancing public engagement of chronic bowel conditions, a short film was made (Gut Reaction) examining the lives of four individuals with different bowel conditions and what scientists and clinicians are doing to help alleviate the pain experienced by such individuals. This film was then screened at a science festival where a pre- and post-film survey was conducted alongside follow up semi-structured interviews with a small subset of those who had expressed willingness to engage in such an interview. Although films have been used for public engagement and health campaigns, there is a lack of a robust evaluation of such methods. As such, there is no knowledge of impacts and outcomes, jeopardising funding of such projects. Overall, the pre- and post-film surveys demonstrated that the film had increased the attendees’ understanding of chronic bowel conditions, how they are treated, what research is on-going and the likelihood of discussing bowel conditions with friends and family. The follow-up interviews were analysed through the constant comparative coding process. The analysis revealed that participants have a strong belief that bowel conditions need to be part of normal conversations, and the understanding of such conditions, and the people who experience them, needs to be improved by society. Our participants hold that this is crucial for people who experience from such conditions, not least to be able to access help sooner and suffer less. Finally, our participants discussed two strategies to achieve this societal openness and tackle the sense of shame around these issues: one involving role models and the other the media. In summary, Gut Reaction appeared to have met its objectives of improving the viewers’ awareness and understanding of chronic bowel conditions, as well as removing some of the stigma and taboo that surround discussions about these conditions.

## Introduction

Normal functioning of the gastrointestinal (GI) tract is often taken for granted. Indeed, many of the GI tract’s physiological functions, such as food digestion and absorption of nutrients, occur without conscious awareness and happen largely under the auspices of the enteric nervous system, a branch of the autonomic nervous system (Greenwood-Van Meerveld et al, 2017). However, conscious sensations such as urgency, discomfort and pain can also arise from the GI tract through the activation of sensory neurones tuned to detect specific stimuli. Although in many cases such sensations pass rapidly, a number of chronic conditions are associated with bowel dysfunction and pain (Hockley et al, 2018), such as inflammatory bowel diseases (IBD, consisting of ulcerative colitis, UC, and Crohn’s disease, CD) and irritable bowel syndrome (IBS). Ideally, preventative treatments and cures would prevent bowel pain occurring in these conditions. However, in the absence of such measures, research is also focused on improving understanding of how sensations arise in the GI tract to enable development of treatments aimed at preventing pain with minimal side effects. This is a particular issue when considering GI tract pain, for example: an individual experiencing pain with IBS with constipation (IBS-C) would not wish to be prescribed an opioid with constipation as a known side effect. One potential route to targeted therapy arises from the recent identification of different sensory neurone populations innervating the distal colon (Hockley et al 2019), where transcriptomic analysis suggests both putative nociceptive (i.e. sensitive to noxious stimuli) and mechanosensory subpopulations of colonic sensory neurones: could targeting nociceptive subpopulations result in treatment with minimal side effects?

Alongside academic and clinical research into pathology of the GI tract, it is important to raise public awareness of such conditions. Bowel habits are not usually the subject of polite conversation, even though each and every one of us is aware of the ins and outs of their daily GI activity. Consequently, under the Pathways to Impact element of a Biological Sciences and Biotechnology Research Council funded grant, we sought to make a short educational film that would bring together individuals with a variety of bowel conditions, scientists and clinicians, and that could be used to highlight the different lived experience of those with disorders of the GI tract and what is being done to help them now and in the future. Videos and Q&A sessions are a recurrent method used to communicate scientific research findings to the public (Frewer and Rowe, 2005). Moreover, communication channels can be used to effectively convey relevant information regarding health issues to the population (Smith-Warner et al, 2001, Singletary and Gapstur, 2001;). Kreps (2003) argues that these messages should include an analysis of the health beliefs, values, and orientations of the intended audience. Furthermore, there is evidence of video being used in health educational interventions as a way of informing patients and audiences in several settings (Latif et al, 2016) including specifically, for bowel conditions (Guts UK, 2019).To that end, researchers at the Department of Pharmacology (University of Cambridge) teamed up with Dragon Light Films. Individuals with different bowel conditions were recruited through the UK charity Bowel and Cancer Research (now Bowel Research UK) to be interviewed about their experiences, alongside interviews with scientists working on GI pain and clinicians treating patients with GI conditions.

The objectives for the development of the film were twofold. Firstly, we wished to improve the viewer’s awareness and understanding of chronic bowel conditions and simultaneously help to remove the stigma and taboo that surround discussing such conditions.

Verne (2004), through a survey of 1014 adults compared the understanding of IBD with other four chronic medical conditions (asthma, coronary heart disease, depression, and diabetes). The results showed a lack of understanding of the condition. Only 1.2% of the respondents thought that irritable bowel syndrome affected more people than did the other 4 chronic conditions, and only 8.6% believed irritable bowel syndrome to be the second leading cause of absenteeism from work or school. Nearly half (44.2%) of the respondents stated that, of the 5 disorders, they knew the least about irritable bowel syndrome.

Patients experiencing bowel conditions are considered to tend to have shame feelings (Casati et al., 2000; Kellett and Gilbert, 2001). Specifically, Casati el at (2000) literature review from previous qualitative and quantitative studies identified eight themes reported by people with IBD: loss of energy, loss of control, body image, isolation and fear, not reaching full potential, feeling dirty, and lack of information from the medical community. In addition, others have shown that chronic illness-related shame presents direct and indirect effects on both psychological health and social relationships (Trindade et al, 2020). As such, our first aim was to create a film that helped reduce this stigma.

Secondly, we aimed to use the film to make the clinical, scientific and treatment process of chronic bowel conditions more accessible to the public. We wanted to assess the film’s impact on the public understanding of several factors associated with the film. To evaluate this, we conducted a short, quantitative survey of viewers attending the film premiere at a university run science festival and the subsequent question and answer session, as well as follow-up qualitative interviews with a smaller subset of the attending group.

Overall, the present study had two aims: firstly, to evaluate the efficacy of the “Gut Reaction” video screening and of a question-and-answer session as a method of informing viewers about chronic bowel conditions, and secondly, to evaluate how the film helped to highlight the stigma or taboo around discussing bowel conditions. Although videos have been used and developed for health campaigns and public engagement projects, there is a lack of a robust evaluation of such methods (Haenssgen, 2019). Without this evaluation, there is no knowledge of impacts and outcomes of public engagement which jeopardises funding of such projects (Haenssgen, 2019).

## Materials and methods

The study followed a mixed methods approach with a triangulation objective. Triangulation seeks convergence and corroboration of results from different methods when applied to studying the same phenomenon (Greene et al, 2007). Quantitative data was obtained through the survey distributed at the screening. After this, qualitative data was obtained through a semi-structured interview format with a smaller subset of the survey participants with a mixture of face-to-face and phone interviews. In the following, we provide methods and results for the two methods sequentially.

### Survey Procedure

The survey was conducted as a pre-post questionnaire. It followed recommendations from (Cohen et al, 2007) to avoid leading, biased, and double-barrelled questions. On entry to the lecture theatre, all attendees were offered a double-sided A5 survey sheet and a pencil (if needed) and asked if they would answer six questions before and the last two sections after the film screening and the question-and-answer session took place. There was a letter box for people to confidentially put their sheets into, when they left the lecture theatre at the end of the event. The survey questions were structured into 3 parts: Part 1 contained pre-impact-oriented questions, part 2 post-impact-oriented questions, and part 3 demographic and contact details to enable follow up interviews where suitable. The full survey is presented in annex I.

The participants in this study were selected randomly from the attendees of the Gut Reaction screening and the question-and-answer session at the 2019 Cambridge Science Festival. Attendees at the screening were not selected in any way: the event was advertised in a variety of media channels (email, website, Twitter etc.), although geographical vicinity is likely to have been an effect limiting ability to attend. The screening took place in a lecture theatre and had 352 attendees. In total, 242 people filled out the feedback survey on the screening of Gut Reaction and the subsequent panel discussion. The survey consisted of two parts – 6 questions before the film section, and another 10 afterwards. While broadly speaking most participants filled out most questions of both surveys, not every participant filled out every question: the lowest response rate to a pre-film survey question was 238; the lowest response rate to a post-film question was 211.

Of the 242 participants, 233 provided their gender and age, 65.2% being female, the average age being 50.7 years with a range of 11 – 85 years of age (Figure 1).

**Figure 1.**
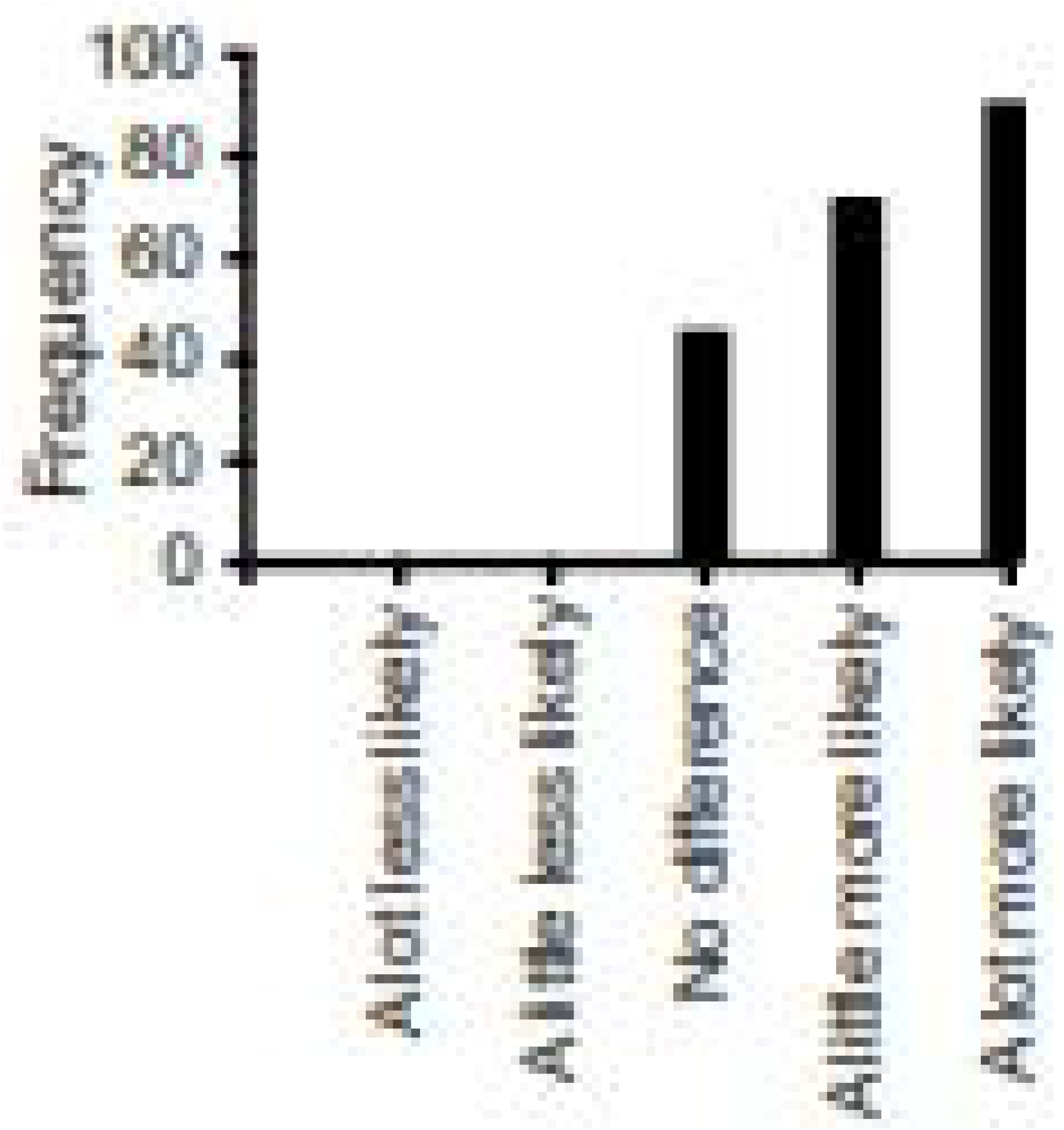
Age range of participants.

**Figure 2.**
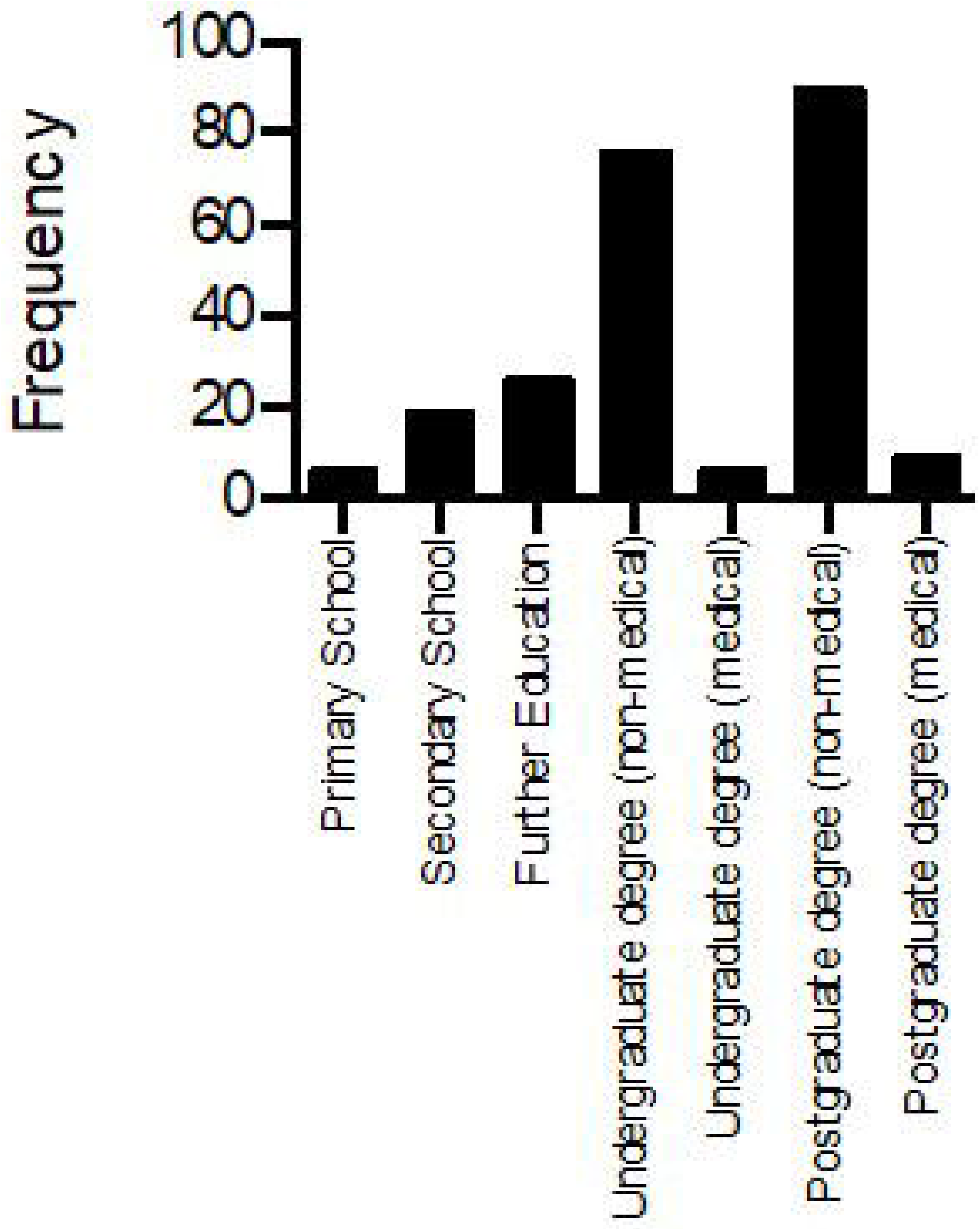
Education attainment of participants.

Of the 232 participants who provided details on their education, 78% had at least an undergraduate degree, and 6.5% had an undergraduate or postgraduate medical qualification, i.e. participants were well educated compared to the general UK population (40.2% of the UK population has a degree according to the Office of National Statistics (2020)). The population of participants that also had a medical education background was greater than in the general UK population (OECD, 2020).

73% of participants stated that they, or someone they knew, lived with a chronic bowel condition. Forty-eight percent stated that they, or someone they knew, had experience of bowel cancer. Of the 183 participants that answered a question relating to their personal experience of regular bowel pain, 19% said that they were currently experiencing regular bowel pain and 24% said that they had experienced regular bowel pain in the past (participants could circle “yes” to both options) and of the 62 participants who answered a question relating to whether or not over-the-counter medications were sufficient to manage the pain, 44% said that they were.

### Survey Results

Data from the questionnaires were analysed using paired Student’s t-tests to examine differences between the questionnaire responses before and after the screening. The results are presented in two sections: The first section describes findings relating to participants’ survey responses, the second section reports the interview findings with the key themes that reflect respondents’ views. Figure 3 a-f. On every parameter assessed, participants reported a statistically significant increase in their knowledge/understanding of all inquired aspects.

**Figure 3.**
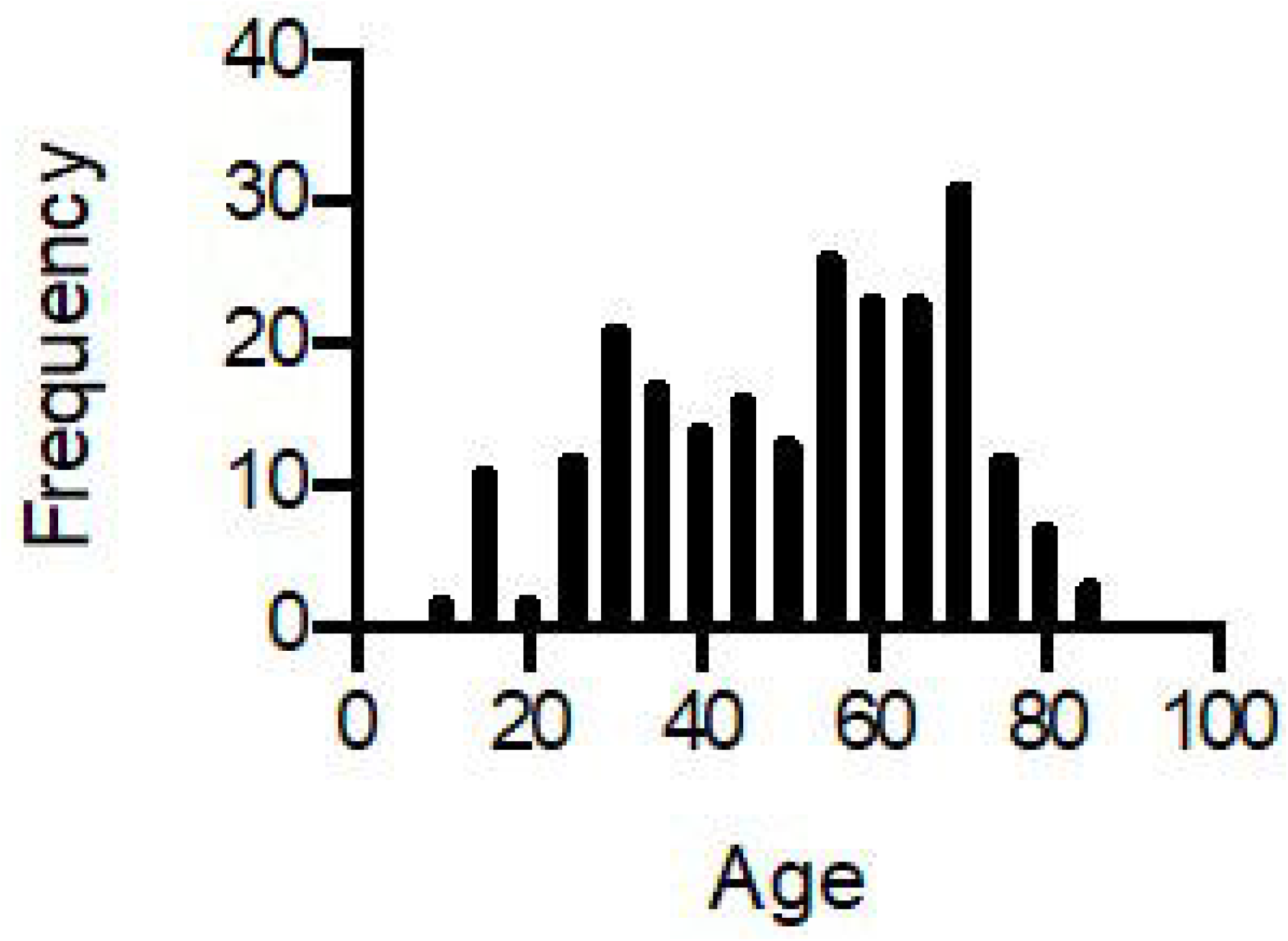
Participant responses to the following identical questions before vs. after viewing “Gut Reaction”. **a)** How much do you know about chronic bowel conditions? **b)** How do you rate your understanding of chronic bowel conditions? **c)** How would you rate your understanding of how a chronic bowel condition can impact your life? **d)** How would you rate your understanding of the treatment of chronic bowel conditions? **e)** How would you rate your knowledge of the main symptoms of chronic bowel conditions? **f)** How would you rate your understanding of how scientists conduct research into pain and chronic bowel conditions? *Each dot on all graphs below represents one participant’s answer and the red lines show the mean ± SEM (*\*\*\*\* *= p* < *0*.*0001 with Student’s paired T test)*

**Figure 4.**
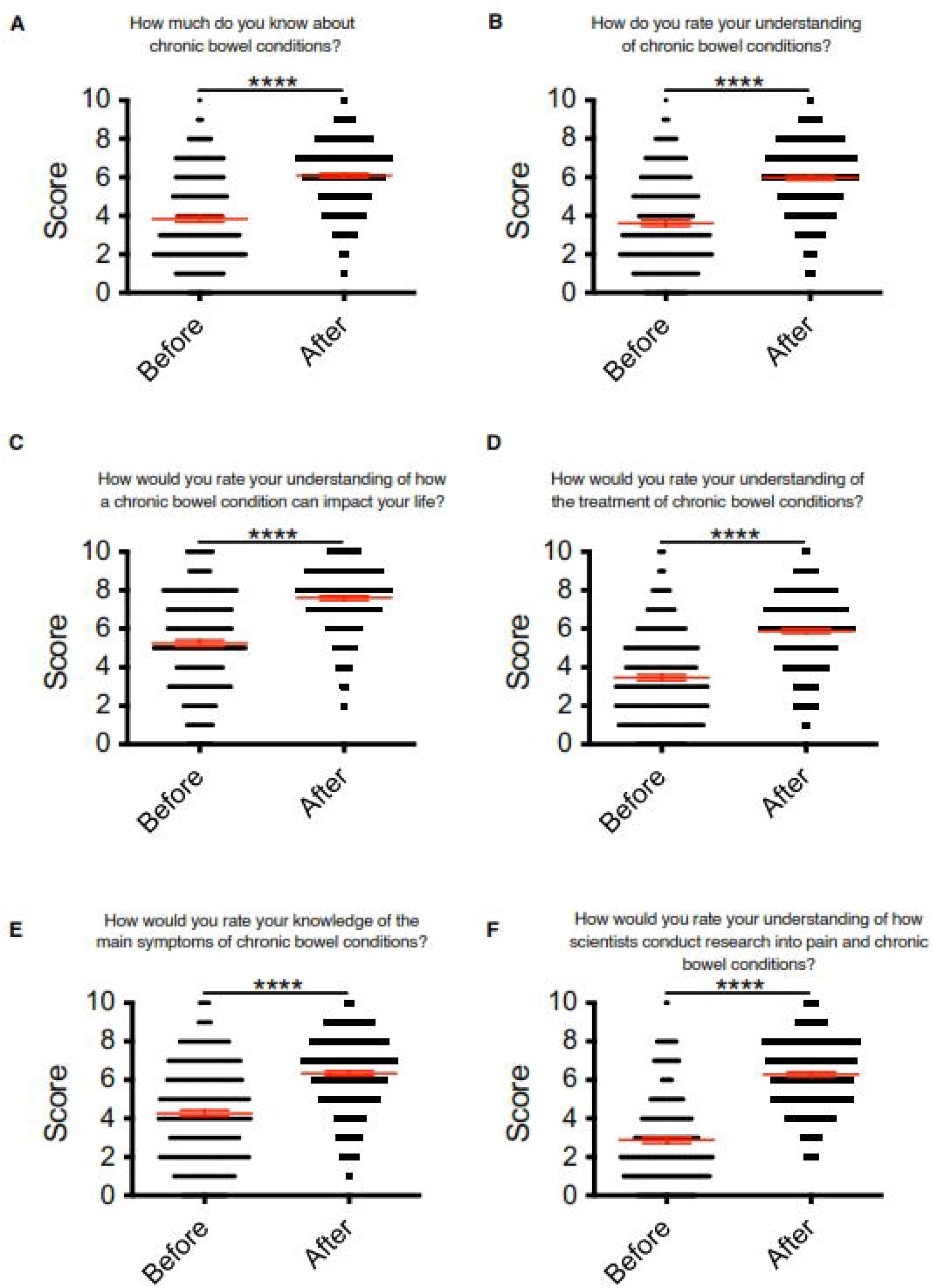
Range of responses to whether a participant was more or less likely to discuss bowel conditions with friends and family because of seeing “Gut Reaction”.

The results show that before the screening of the film, on average, participants rated their understanding closer to poor (below 5/10) than to excellent, on the Lickert scale, for most questions. Participants self-reported that they had limited understanding of chronic bowel conditions, of treatments, symptoms and of how research is conducted into these conditions.

Previous research, like Verne’s (2004) survey, has found that there is a lack of public knowledge in relation to bowel conditions. Interestingly, even with our self-selected audience – audience that booked to attend a talk on bowel conditions at a science festival – the level of understanding was limited. The only question in which participants on average answered above 5 was the one that asked participants if they had an understanding on how these conditions impact one’s life. The higher score in this question is likely to be related with the presence in the audience of individuals living with chronic bowel conditions, or family members of such individuals.

Findings like the one revealed by Verne (2004) highlight the need for public education initiatives to raise awareness and knowledge about the prevalence and impact of bowel conditions. This was one of the objectives of the Gut Reaction film as previously described. The results of the pre and post survey demonstrate that self-reported understanding of the different aspects related to bowel conditions increased significantly in all aspects. These results offer strong evidence for the importance of films such as Gut Reaction in public engagement initiatives, which as stated in needed (Haenssgen, 2019).

The survey results also indicated that the audience had a keen interest in bowel conditions, such that 74.55 said that they had discussed a bowel condition with friends and family. Moreover, 77.3% said they were more likely to discuss these conditions after watching Gut Reaction. This is a strong indicator that the film will potentially generate discussion in relation to bowel conditions, which is essential to diminishing stigma (Trindade, 2020). Finally, the panel discussion held after the screening of the film generally received positive reviews, scoring an average of 6.6 on a scale of 0 (no improvement) to 10 (definite improvement) for understanding how the bowel works and what can go wrong. Lastly, when asked if participants would recommend Gut Reaction to friends, it scored an average of 7.7 on a scale of 0 (not recommend) to 10 (definitely recommend).

### Follow-Up Interviews

Seventy-one out of the 242 survey participants stated they were happy to be contacted for an interview. Forty-four were subsequently contacted (some being excluded for being under 18 years of age or failure to establish contact with others due to illegibility of handwriting and/or emails bouncing back). Of these, 11 replied and were interviewed. Table 1 describes the interviewees’ characteristics.

**Table 1.**
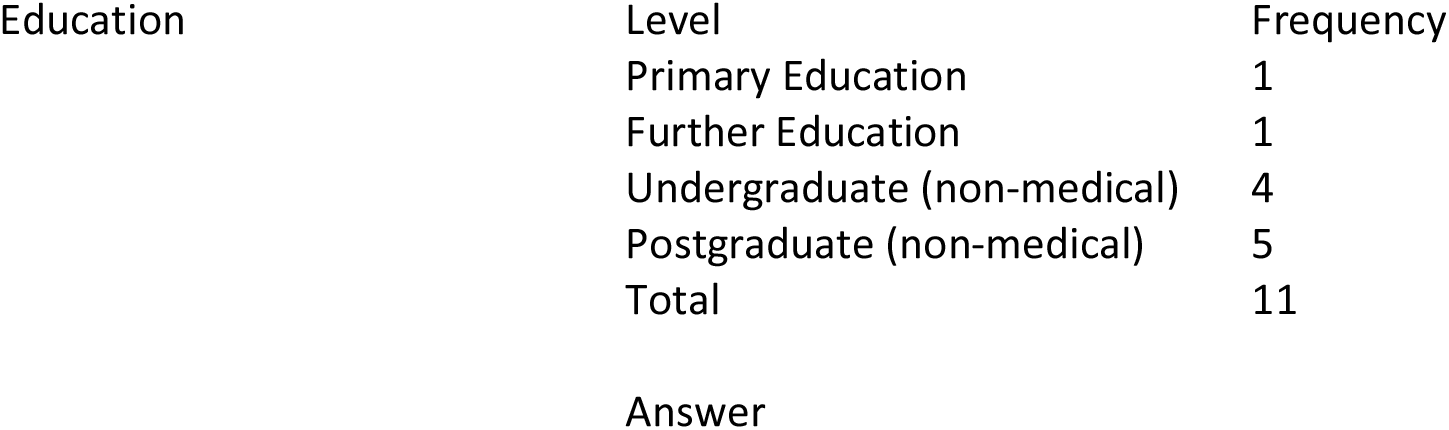
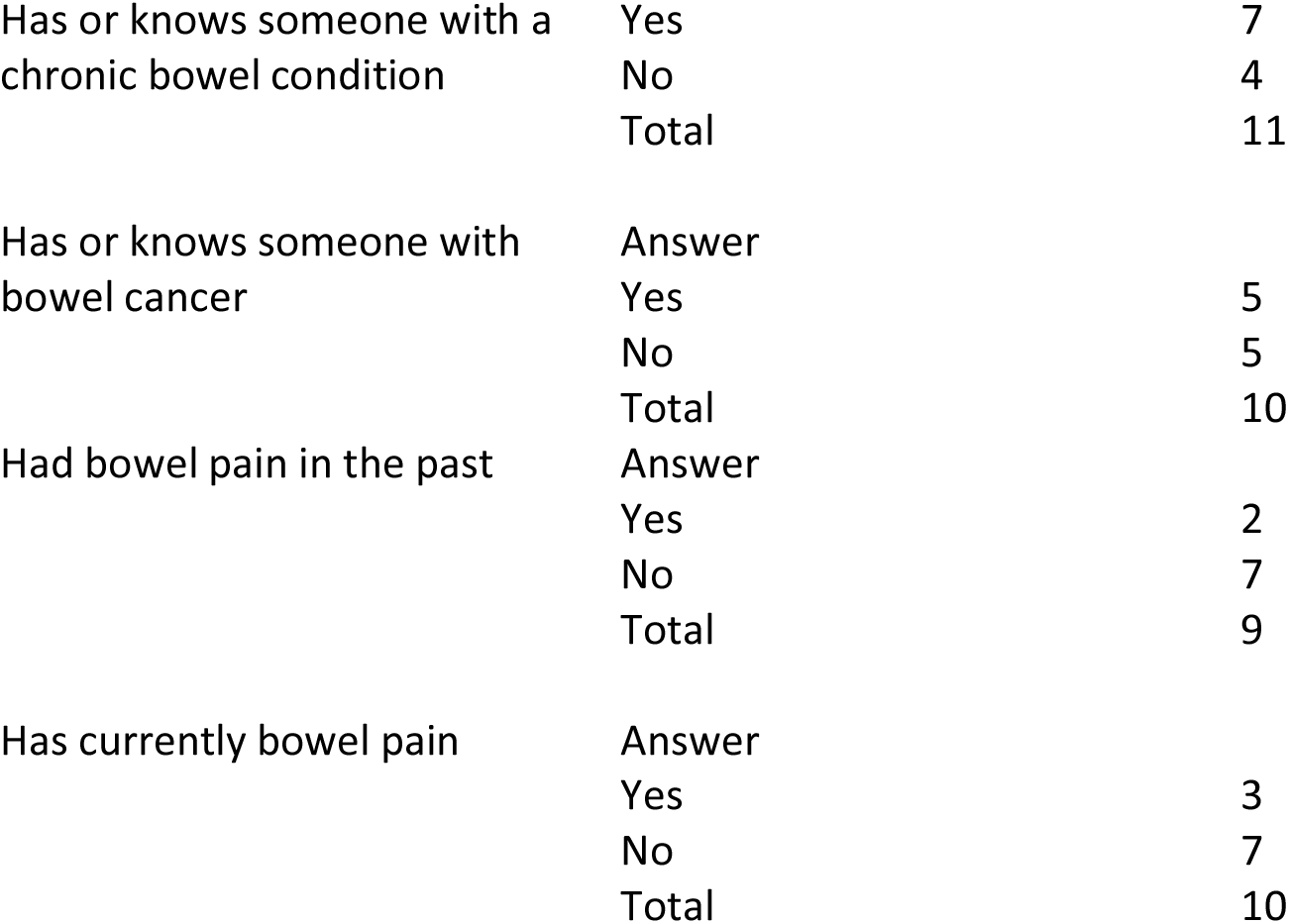
Interviewees’ characteristics.

Participants were asked if they had talked about bowel issues in the months following the screening. Eighty percent of the interviewees said that they did. This offers even further evidence of the impact of the video. In the survey, immediately following the screening of the video, we asked participants if they were more likely to talk about bowel issues to family and friends. Seventy-seven percent said they were. In the subsample we interviewed, we obtained a higher percentage (80%) of participants who reported having done this. These results highlight the potential of videos such as Gut Reaction in generating discussion about sensitive topics.

The interview recordings were transcribed and anonymised. The qualitative analysis of the interview transcripts borrowed principles from the constant comparative coding process. In the process of coding, three concepts were used: codes, categories, and themes. Green (2008, p. 71) asserts that “novice researchers typically make few distinctions among analysing for themes, categories, and codes”. Therefore, in this study special attention was given to the differentiation between the three concepts. The analysis was facilitated by the qualitative computer software package NVivo.

The coding and sorting of the interview data resulted in key themes that reflect respondents’ views. The coding started with abstracting topics from the transcripts. This was achieved in two sequential ways. Firstly, the interviews were manually transcribed to MS Word files with the help of Potplayer software. Next, the researcher identified the initial concepts the participants were discussing. Secondly, the initial coding continued with NVivo 12 which enabled the grouping of related concepts. In NVivo, these related concepts are organised in containers named nodes. These initial nodes are topics, ideas or abstractions that come from the study (Bryman, 2008). At this point, there were 15 initial codes. Different methods available in NVivo were used to reflect on the codes that were developing: Firstly, the code files were analysed for consistency and frequency. Secondly, the framework matrices were created to analyse the codes across all participants. The framework matrices enabled analysis of initial nodes. Further analysis was carried out to attain the seven higher-order categories. In the following stage, the categories were merged and re-named into four main themes: “Comparison with how other diseases became openly talked about”, “The need for openness and understanding”, “Suffered more than needed”, “Strategies: role models and the role of media”.

The themes were tested for reliability. A separate researcher was consulted to ensure agreement. After this consultation, both intra- and intercoder agreement was above 90%. This is above the range defined by Miles and Huberman (1994).

In the following, we focus on four key themes discussed by the interviewees: comparison with other diseases; openness and understanding; unnecessary suffering; and communication strategies.

## Interview Results

### Comparison with how other diseases became openly talked about

When commenting on the film’s importance, many participants drew comparisons with other diseases and how those are now discussed commonly and openly.

For instance, Joanne stated:

> “Things like cancer have come a long away. Ten, 15 years ago people didn’t talk about it. Now it is different. Somehow that’s the point we need to get to get with this.”

Joanne’s comparison is reflective of the work done to remove the taboo around, and open conversations about, cancer by charities and governments alike, including through online and offline campaigns (e.g. McMillan, 2020). Several interviewees made the same comparison to cancer in particular:

Jane stated:

> “People know a lot about cancer but something like this [IBS] can have a lot of impact on people’s life”.

Andrew stated:

> “Because you see it with prostate cancer. If you go back around 10 years, no one heard of it”.

It was clear that our participants recognised the value of diseases being openly talked about and being known. The impact that health communication can have in early detection, prevention, diagnosis and treatment has long been recognised (e.g., Kreps & Sivaram, 2008). Openness and conversation with families and friends plays a particular role to increase the impact of such health communication (e.g., Street, et al, 2009).

### The need for openness and understanding

The next theme focuses on reasons why participants think that openness is not always present.

John suggested:

> “People don’t want to discuss faecal matters at dinner table which I can understand but there are many people that only meet at the dinner table, it’s one of the great meeting places. It should be more understood”.

Indeed, multiple interviewees recognised that they found it difficult to talk about these issues all the while they also believed people needed to be more open about and aware of the subject. Previous research has shown that it is detrimental not to discuss these issues. For instance, Palmer et al (2014), concluded in their qualitative study that open discussions should be designed and evaluated in order to increase the uptake of bowel cancer screening:

> “Participants (…) described being influenced by discussions with family members, friends, and health professionals. (…) They also recalled supportive discussions in which their concerns about or aversions (…) were discussed and challenged. (…). In addition, they reported that becoming aware that a family member or friend had developed bowel cancer influenced them to take part”. (p. 1709)

One of our interviewees also mentioned why it is important to discuss the subject more and to make it a normal topic of conversation.

Amy stated:

> “(I) think it’s important to get the general public view because there can be stigma against it and people can be quite shy if they have the disease”.

To increase the level of comfortableness and willingness to discuss bowel issues, it is important to normalise the subject within society. As outlined by Thompson (2013), there are deeply ingrained definitions of faeces as a taboo substance and rigid social rules surrounding how we should appropriately deal with them in society. The Guts UK charity website (2020) tries to establish the counter-narrative that “[i]t is ok to talk about poo! And it is really important that people don’t suffer, or worse – die – of embarrassment, when it comes to talking about their bowel habits.”

### Unnecessary suffering

In the third theme, our participants discussed the perils posed by embarrassment.

Jane stated that it was “important to change it because I personally suffered longer than I should have done because I was too embarrassed to talk with anybody, even go to the doctor about it. It needs to be seen as another illness people have and there is nothing to be embarrassed about so people can go and get help and not suffer.”

Jane gives her personal account of how feeling shame led her to not being open about her condition and as a result to suffer more. She mentions how gut issues being seen as embarrassing and not talked about as other diseases are makes the lives of those with such conditions more difficult. These feelings of shame are common on patients with bowel conditions, as reported by Hall et al. (2005). Furthermore, Walker et al. (2008) have described patients with IBD being at least twice as likely to develop a depressive disorder in comparison with normal controls with similar ages and backgrounds.

Jane was not the only one reporting these feelings, Megan also argued in a similar way:

> “I think it is really important because it helps to get help sooner, helps scientists understand it better, if you suffer from it you feel more supported and less alienated”.

Megan thus highlighted three key points: that not feeling shame allows you: 1) to obtain assistance sooner and 2) to feel supported; and 3) how important it is for researchers to hear from sufferers.

What our participants disclose here is supported by Trindade et al. (2020) on their quantitative research with IBD patients. Patients react to disease related shame by forming avoidance patterns. These patterns then jeopardise the patient’s life amplifying the damaging effect of the disease. As such, the avoidance patterns’ possible impacts are twofold. They may lead to possible sufferers not discussing their issues, which leads to disease not being tackled promptly; and, they may compromise important areas of the patient’s life, inhibiting the engagement in actual valued activities and therefore damaging their psychological health and social relationships.

### Strategies: role models and the role of media

In the last theme discussed by our participants, they mentioned strategies that could help bowel issues become less of a taboo and a more open topic.

Research participants shared their views on what could be beneficial in making discussions about bowel issues more open and how to overcome the sense of shame. One approach that participants mentioned were role models. The Gut Reaction video had testimonials from a diverse range of individuals with different bowel conditions and our research participants highlighted that fact. For instance, Megan stated:

> “it actually showed kids, and young people, older people and also different problems they have, the different problems can have and impact. by having that breadth of people it had a bigger impact”

And Rob mentioned:

> “what it means to them in their everyday life and all four of them were just so honest and that is quite striking to be talking about something personal”

Our participants are highlighting two key aspects of good communication messages, narrative and role models. Previous research has shown that when people are transported into a narrative world, they are more likely to change their behaviour (Green, 2006). Our participants reported how involved they became in the narrative and the impact our sufferer’s narrative had on them. One way in which viewers are transported through the narrative is by creating connections with characters (Green & Brock, 2000). If the viewer likes or identifies with a character, seeing them as a role model, the implications of events experienced or assertions made by the character will more likely have an impact on the viewer (Green, 2006). This is exactly what our participants are describing when discussing the characters on gut reaction. Furthermore, as identified by Ruth, the fact that film had a diverse range of people represented who shared their stories increased its potential as a role model narrative. The film’s use of first-person narratives aligns with previous research that suggests that narrative communication is intrinsically more persuasive than didactic communication; information communicated through narratives often results in greater acceptance through narratives’ ease of processing and comprehension (Graesser, Olde, & Klettke, 2002; Schank & Abelson, 1995). Throughout the film, we hear the narratives of a diverse group of individuals with different chronic bowel conditions. This is also aligned with previous research that has shown that first-person narratives increased experience-taking and altered behaviour more than third-person narratives (Kaufman & Libby, 2012) and that first-person narratives resulted in greater identification, which itself mediated attitude change as compared to third-person narratives (de Graaf, Hoeken, Sanders, & Beentjes, 2012). Likewise, a review of studies within the field of health communication suggests that first-person narratives were more influential than third-person narratives in health decisions (Winterbottom et al., 2008).

Finally, the interviewees also argued for ways in which discussion over bowel issues could increase. A recurrent view shared was the role of mass media.

Megan mentioned:

> “short segments that come across on tv, broad media is very helpful and if it is presented that way many people would have seen it, it can spark discussion.“

And Dawn and Rob also argued:

> “social media would be quite useful, it’s getting things in the modern society, if people with Chron’s are shown on tv it becomes more society awareness”
>
> “TV programmes could be a way, posters on public toilets”

Our participants mentioned different platforms that could be used, but often focused on traditional media and especially TV. In a time of social media and streaming platforms, mainstream TV may seem of secondary importance when it comes to health messages, but our participants highlighted its continued relevance. Indeed, in the “public attitudes to science” report (DBEIS, 2019), 47% of evaluated UK citizens state that TV is still the main source of information about science. Furthermore, the use of TV and mass media for health messages is well documented but its efficacy is mixed (Randolph et all 2004). In particular, Randolph & Viswanath (2012) have identified components of successful public health campaigns to include: 1) successfully manipulating the information environment by campaign sponsors to ensure sufficient exposure of the audience to the campaigns’ messages, 2) using social marketing tools to create appropriate messages for distribution and, where possible, message theory and tailoring, 3) creating concomitant structural conditions such as a supportive environment/opportunity structure that allows the target audience to make the recommended change, and 4) understanding the determinants of health behaviour that could potentially lead to desired health outcomes (theory based campaigns). As demonstrated by our findings, the use of role models and well-constructed narratives may similarly contribute to better outcomes and lasting impact at a smaller scale.

## Discussion

This study had two aims: evaluate the efficacy of the Gut Reaction film screening and question-and-answer session as a method of informing viewers about chronic bowel conditions, and secondly, to understand the stigma or taboo around discussing bowel conditions and to explore the barriers to remove them. To achieve this, quantitative and qualitative data was collected under a mixed-methods approach.

In relation to the first aim, the pre- and post-film surveys’ results show a significant increase of the participants’ understanding of bowel conditions, the impact these conditions have on individuals’ lives, but also increased the understanding of the treatments available, of the symptoms of bowel conditions, and of research being conducted on these conditions. Furthermore, in follow-up interviews, we asked survey participants if they had discussed bowel issues in the months following the screening and 80% of the interviewees said that they did. Additionally, some participants mentioned they specifically discussed the film with friends or family months after having watched it. These results showcase the impact informative videos can have on increasing audience understanding. Viewers discussing it with friends and family can multiply this impact.

In relation to the second aim, the semi-structured interviews provided insights into the participants’ views. Subsequent analysis revealed that participants have a strong belief that bowel conditions need to be part of normal conversations, and that society needs to understand better these diseases as well as the people suffering from them. Participants compared this to similar paths, other diseases had gone through in terms of societal recognition. Our participants believe that the same needs to happen with bowel diseases. Our participants hold that this is crucial for people who suffer from such conditions, not least to be able to access help sooner and suffer less.

Our participants discussed two strategies to achieve this societal openness and tackle the sense of shame around these issues: one involving role models and the other the media. Firstly, the participants in this study stressed how the diverse people represented in the film and their stories made an impact. This aligns with research on the importance of identification for comprehension, acceptance, and altered behaviour.

Secondly, our participants debated the media’s importance for the discussion over bowel issues. Participants argued that bowel issues being visible in everyday media would be crucial for their normalization. Mass media health campaigns are already a major tool for public health practitioners, with mixed success. When participants mention how diverse role models sharing strong narratives leaves an impact, they are highlighting the importance of message theory and tailoring. Our participants are identifying what makes the message successful and offering insight on what should be a theory-based campaign. Furthermore, participants mentioned time and time again the importance of changing the wider society environment to make bowel issues more acceptable. Participants emphasised what Randolph et al (2012) describe as the opportunity structure. A successful broad media campaign around bowel conditions needs also to be about changing the wider environment to make it more open acceptable.

In summary, the film Gut Reaction has provided important insights that can be used by others when planning smaller-scale media campaigns around bowel conditions. Moreover, it highlights the importance of providing funding in research projects for impact activities. They can give voice to participants, enable understanding of their views and concerns, and consequently lead to the development of new projects and campaigns based on the learnings obtained.

## Acknowledgements

We are grateful to all those who participated in making the film, Bowel and Cancer Research (now Bowel Research UK) who supported film development and all members of the audience who participated in the survey and follow-up interviews. We acknowledge support from clinicians who took part in the film. We further acknowledge thanks to those who took part in the question and answer session at Gut Reaction’s screening alongside EStJS and JRFH, namely, Deborah Gilbert (Bowel and Cancer Research CEO), Dr Dervila Glynn (Cambridge Neuroscience) and Robert Heuschkel (Consultant Paediatric Gastroenterologist). We thank Dragonlight films (https://dragonlightfilms.com/) for creation on the film. This work was support by a BBSRC grant to EStJS and JRFH (BB/R006210/1).

