## Supplementary material for "Gut Reaction: The Impact of a Film on Public Understanding of Gastrointestinal Conditions": Table 1

| Education | Level | Frequency |
| --- | --- | --- |
|  | Primary Education | 1 |
|  | Further Education | 1 |
|  | Undergraduate (non-medical) | 4 |
|  | Postgraduate (non-medical) | 5 |
|  | Total | 11 |
| Has or knows someone with a chronic bowel condition | Answer |  |
|  | Yes | 7 |
|  | No | 4 |
|  | Total | 11 |
| Has or knows someone with bowel cancer | Answer |  |
|  | Yes | 5 |
|  | No | 5 |
|  | Total | 10 |
| Had bowel pain in the past | Answer |  |
|  | Yes | 2 |
|  | No | 7 |
|  | Total | 9 |
| Has currently bowel pain | Answer |  |
|  | Yes | 3 |
|  | No | 7 |
|  | Total | 10 |

Table 1 - interviewees’ characteristics.
